# Stem Cell Plasticity and Niche Dynamics in Cancer Progression

**DOI:** 10.1101/056762

**Authors:** Noemi Picco, Robert A. Gatenby, Alexander R. A. Anderson

**Affiliations:** N. Picco is with the Integrated Mathematical Oncology Department, H. Lee Moffitt Cancer Center and Research Institute, Tampa, FL, USA and the Wolfson Centre for Mathematical Biology, Mathematical Institute, University of Oxford, UK.; R. A. Gatenby is with the Department of Radiology and the Integrated Mathematical Oncology Department, H. Lee Moffitt Cancer Center and Research Institute.; A. R. A. Anderson is with the Integrated Mathematical Oncology Department, H. Lee Moffitt Cancer Center and Research Institute.

**Keywords:** Cancer stem cells, ductal carcinoma in situ, hybrid discrete-continuum cellular automata, niche, plasticity, stemness, tumor microenvironment

## Abstract

**Objective:** Cancer stem cells (CSCs) have been hypothesized to initiate and drive tumor growth and recurrence due to their self-renewal ability. If correct, this hypothesis implies that successful therapy must focus primarily on eradication of this CSC fraction. However, recent evidence suggests stemness is niche dependent and may represent one of many phenotypic states that can be accessed by many cancer genotypes when presented with specific environmental cues. A better understanding of the relationship of stemness to niche-related phenotypic plasticity could lead to alternative treatment strategies.

**Methods:** Here we investigate the role of environmental context in the expression of stem-like cell properties through in-silico simulation of ductal carcinoma. We develop a two-dimensional hybrid discrete-continuum cellular automata model to describe the single cell scale dynamics of multi-cellular tissue formation. Through a suite of simulations we investigate interactions between a phenotypically heterogeneous cancer cell population and a dynamic environment.

**Results:** We generate homeostatic ductal structures that consist of a mixture of stem and differentiated cells governed by both intracellular and environmental dynamics. We demonstrate that a wide spectrum of tumor-like histologies can result from these structures by varying microenvironmental parameters.

**Conclusion:** Niche driven phenotypic plasticity offers a simple first-principle explanation for the diverse ductal structures observed in histological sections from breast cancer.

**Significance:** Conventional models of carcinogenesis largely focus on mutational events. We demonstrate that variations in the environmental niche can produce intraductal cancers independent of genetic changes in the resident cells. Therapies targeting the microenvironmental niche, may offer an alternative cancer prevention strategy.

## I. Introduction

THE cancer stem cell (CSC) hypothesis proposes that cancers arise from a small population of cells that, similar to somatic stem cells, are self-renewing and give rise to subpopulations of more differentiated cells with limited capacity for proliferation [1]. Cancer stem cells are thought to have a slow cell division cycle, active DNA repair system, and, most importantly, are resistant to conventional therapies, causing disease relapse after treatment. In this paradigm, tumors are assumed to contain cells at various stages of differentiation, from stem-like cells, providing the pool of self-renewing cells, to the terminally differentiated ones, with limited proliferative potential. Markers of stemness have been identified for almost all cancer types but interestingly not all markers agree [2], [3].

In Fig. 1, pathological samples from three different breast cancer patients are shown. The variation in structural tissue organization across patients can be easily appreciated. The first sample [Fig. 1(a)] demonstrates well-defined ductal-like structures with a hollow lumen. In the second sample [Fig. 1(b)] similar ductal structures can be recognized, although cells have lost their polarization and structural organization is lost as proliferating cells fill the lumen. This morphology is typical of the *Ductal Carcinoma In Situ* (DCIS). In the last pathological slice [Fig. 1(c)], the ductal structure is completely lost giving way to structural disorganization, indicating loss of differentiation.

**Fig. 1.**
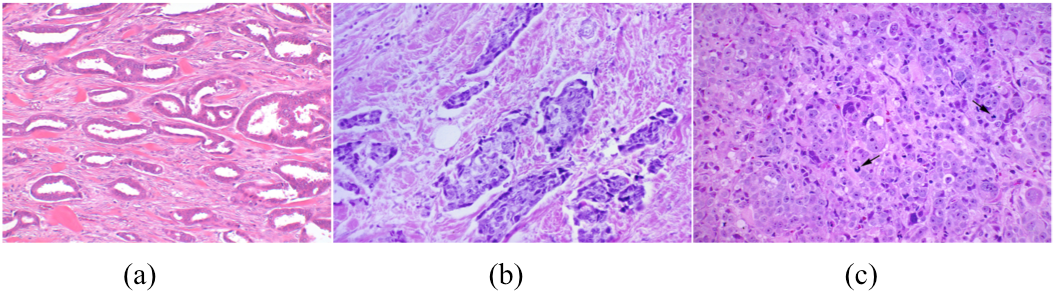
Histology of Breast cancer at different stages of progression. (a) Well differentiated tissue, showing well defined ductal-like structures composed of tumor cells (darker pink) and hollow lumen (in white). (b) Moderately differentiated tissue, ductal-like structures are still clearly defined, but without any lumen as they are filled with tumor cells (darker stain). (c) Poorly differentiated tissue, the ductal structure is completely lost, only a dense field of tumor cells is observed.

DCIS is thought to follow a temporal progression from well-differentiated ductal organization [as in Fig. 1(a)] through a moderately differentiated one [as in Fig. 1(b)] to a poorly organized and highly invasive cancer [as in Fig. 1(c)]. This progression of pathological stages is often described as “somatic evolution” and is conventionally viewed as a process driven solely by accumulating mutations.

The role of CSCs in the evolution of breast cancer remains unclear. The hierarchical model proposes that only a fraction of cancer cells are CSCs with the ability to self-renew indefinitely [4]. In this model, most cancer stem cells are passing through differentiated states, similar to the development of normal tissue. These cells have limited proliferative capacity and are, thus, unable to recapitulate the tumor if the CSCs are lost. Therefore, in this model eliminating CSCs will effectively eradicate the tumor. An alternative model proposes that stemness is a terminal phenotypic state that can be achieved by any cancer cell [4]. This implies that most and perhaps all cancer cells can adopt stem-like properties with appropriate environmental cues in a unidirectional manner.

Recently, a third hypothesis has been proposed: that stemness is merely one component of the reaction norm of a cancer cell. That is, it represents one of many phenotypic states that can be expressed by the same cancer genotype depending on environmental conditions – similar to, for example, variations in the phenotype of a tree during summer or winter. Thus, “sternness” can be gained and lost by each cancer cell over time depending on local environmental conditions [5], [6]. However, the precise mechanisms behind the interconversion between CSC and non-stem cancer cells are still largely unknown.

Here we investigate one possible mechanism of niche-modulated stemness by mathematically framing the hypothesis that CSCs represent a transient phenotypic state governed by interactions with local environmental conditions. Our model preserves the hierarchical organization inherent in the two other paradigms, however, it permits continuous reprogramming of cell state by environmental cues.

Our work builds on a number of previous computational investigations of CSC dynamics (for an extensive review, see [7]). Cancer stem cell plasticity has also been previously modeled as dedifferentiation of progenitor cells, thus relaxing the conventional unidirectionality of the differentiation process [8] – [11]. However, in the CSC modeling community little emphasis has been put on the drivers (we argue, environmental) that modulate stem cell plasticity [12]. Here we develop a mathematical model of context-driven cancer stem cell plasticity in which stemness continuously varies across a phenotypic spectrum, directly modulated by environmental cues.

## II. The Microenvironment: A Modulator of Stemness

In normal somatic stem cells the microenvironment is a well accepted regulator of stemness through the stem cell niche [13]. Consisting of factors such as ECM, growth factors and metabolites, this niche is also important in cancer [14]. The tumor microenvironment is already an accepted major modulator of the stemness phenotype in a variety of cancers [15], [16]. According to the CSC hypothesis, cancers arise from cells with embryonic\stem resemblance whose malignant phenotype is triggered when located in an abnormal environment, the *cancer stem cell niche* [17]. The broad definition of niche as the permissive and supportive environment for cancer stem cells is derived from its analogue in normal somatic stem cells. Thus the niche is considered to be the sum of factors that constitute the tumor environment, promoting the induction and maintenance of stem-like properties in cancer cells, and protecting them against treatment toxicity.

The extracellular matrix (ECM) is a major component of the tumor microenvironment and exerts a primary function in many processes of cell biology, including cell differentiation. Indeed, the role of ECM is not only limited to the structural support (scaffolding), but is also involved in signal transduction, determining cells fate [18]. For instance, by probing the ECM, cells can sense changes in the microenvironment and respond accordingly initiating a cascade of signaling events. Most importantly, ECM anchorage determines the balance between self-renewal and differentiation in stem cell populations, by controlling cell polarity that in turn determines symmetric or asymmetric division [19].

The ECM, together with stromal cells, a number of soluble factors, signaling pathways and environmental conditions have been proposed to be part of the cancer stem cell niche. Endothelial cells have also been shown to induce and maintain the self-renewal ability of glioma and breast cancer stem cells (activating the Notch pathway [16]) and to enhance their resistance to radiotherapy [20]–[22]. In colon cancer the balance between differentiation-inducing and stemness-promoting factors (i.e. Wnt signaling and BMP signaling, respectively) is dependent on the niche, dictating the malignancy of the tumor [23]. Specifically, it has been observed that myofibroblasts play a key role in the reactivation of the Wnt pathway, driving differentiated tumor cells towards a more stem-like phenotype. TGF-β and other EMT-inducing factors not only regulate cell stemness but also stimulate the proliferation of the cancer stem cell pool, thus enhancing invasiveness and metastasis of breast cancer [24]. However, recent work has established that hybrid states, when cells are between epithelial and mesenchymal, are where stemlike phenotypes are more likely to emerge [25], [26]. Therefore, the relationship between EMT-inducing factors and stemness is more nuanced and beyond the scope of this work.

In addition to these cellular and soluble microenvironment components, a number of stress factors affecting the environment itself have been identified as drivers of dedifferentiation. Among these are hypoxia [27] and inflammation caused by cell death events [28]. Neuroblastoma and small-cell lung carcinoma cells exposed to hypoxia and oxidative stress have shown enhanced migratory, invasive and tumorigenic ability [29]. Additionally, hypoxia induced factors (HIFs) have been shown to induce upregulation of genes linked to stemness properties, such as invasion, treatment resistance, and self-renewal [30]. A separate study showed that acidosis, regardless of oxygen availability, causes a similar promotion of stemness [31]. Importantly, it has also been reported that stimuli following cell death events, such as apoptosis and therapy-induced necrosis stimulate the tumorigenic potential of the cells in the surrounding environment, promoting the dedifferentiation process. Specifically, the same proteases responsible for apoptotic cell death have been observed to induce growth signal and tissue regeneration, constituting the so-called *phoenix rising* effect [32]. Finally, radiation has been observed to stimulate the emergence of a CSC fraction in breast cancer, able to repopulate the tumor [33].

The complex interplay between the tumor cell population and the niche makes it difficult to disentangle just who is driving who. We believe that there is now sufficient evidence to suggest that the hierarchical structure of a tumor is constantly redefined by interactions between the tumor and its environment. The niche can dictate the degree of cell stemness, driving it to a more differentiated phenotype or maintaining its stem cell fate. Conversely tumor cells can have a determining role on the niche, for example expressing signals that in turn limit or enhance the cancer stem cell pool.

Given the complex network of environmental signals and factors mentioned above, we will describe the role of the environment in modulating stem cell plasticity by considering a generic *dedifferentiation signal.* This signal will be representative of the sum of multiple factors described above, which when combined with ECM density will determine stem cell phenotypes.

## III. Multiscale Hybrid Discrete-Continuum Model

In this paper we explore the hypothesis that all cancer cells possess the ability to acquire an invasive stem-like phenotype, capable of initiating or repopulating a tumor. In our model, stemness has two main characteristics. First, it can vary in grades on a scale that spans from the most stem-like phenotype (e.g. highly invasive, with self-renewal ability), to a fully differentiated one (e.g. with exhausted proliferative ability and poorly invasive). Secondly, this stemness phenotype is plastic -directly modulated by the environment. The cancer stem cell niche, namely an environment favorable to stemness, is necessary to maintain cancer cells at a highly stem-like phenotype. Conversely, loss of these favorable conditions will produce gradually more differentiated, less stem-like progeny, with reduced invasive ability and limited proliferative potential.

We examine the role of stem cell plasticity through the lens of early ductal carcinoma by implementing a hybrid discrete-continuum (HDC) model. This multiscale hybrid modeling framework, first introduced by Anderson [34]-[36], allows for the description of discrete individual cell events coupled with continuous environmental factors.

Stem cell plasticity is modeled by considering a continuous gradation of stem like phenotypes, from true stem to truly differentiated. Each cell is characterized by a *degree of differentiation D^i^*, taking the minimum value *D*^0^ for a fully stem-like cancer cell, and the highest value *D^N^* for a fully differentiated cancer cell, where *N* is the total number of degrees (or grades) in the spectrum of stemness. A cancer cell at a given *D^i^* possesses three phenotypic traits that characterize its level of stemness, as illustrated in Fig. 2. The traits are proliferation potential, ECM degradation and production rates. So a true stem like cell will have maximum ECM degradation, and proliferation potential. Whereas a fully differentiated cell will produce ECM, and not proliferate.

**Fig. 2.**
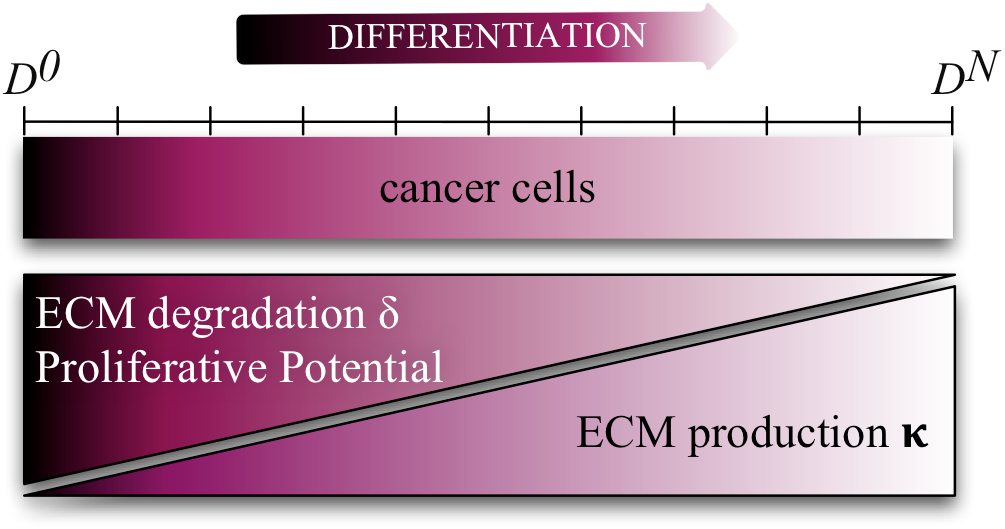
Definition of cancer cell phenotypes, according to their grade of stemness. Each cell is characterized by a degree of differentiation *D^i^*, varying from *D*^0^ (highest stemness) to *D^N^* (fully differentiated, poorly stem). Corresponding to each *D^i^* is a specific phenotype that determines the cell’s ability to remodel the ECM (production and degradation rate) and its proliferative potential. The variation of parameter magnitude, as a function of stemness degree, is represented below the stemness scale.

Plasticity is then simply a cancer cells ability to shift back and forth on the scale of differentiation, changing its *D^i^* according to the environmental conditions, i.e. the niche. Fundamentally this means that a fully stem-like cancer cell can become terminally differentiated (*D^N^*) and subsequently return to a full stem state (*D*^0^) given the right cues.

For simplicity we will focus on the interactions between cancer cells and two environmental factors modulating the niche. The first one is extracellular matrix (ECM) that here represents several components of the stromal environment both sustaining tumor proliferation and invasion (e.g. growth factors, scaffold of ECM proteins) and acting as an antagonist (e.g. stromal and immune cells competing for the same resources as cancer cells and limiting their growth, ECM build-up limiting invasion) in a density dependent manner. The ECM density is modeled as a continuous function of time and space *f* = *f*(*x,t*). ECM is both degraded by cell-produced enzymes such as metalloproteinases, and remodeled, via cell secretion of extracellular components, as previously modeled in [37], [38].

The second microenvironmental variable is a *dedifferentiation signal* (DS). Given the complexity of the interactions between tumor cells and environment, and the so far poorly understood mechanism governing the dynamics of the stem-like population, we assume this signal integrates all possible factors stimulating and maintaining the stemness in the cancer cell population. As we discussed above experimental evidence suggests, stemness is primarily induced and maintained by stress factors, and following cell death events. We model DS as a continuous function of time and space *c* = *c*(*x,t*), and assume it is locally increased by cell death events, diffusing throughout the tissue, and decaying at a constant rate. This DS may persist in the system for a time *T_P_*, beyond instantaneous cell death, so we name this parameter DS *persistence*.

The evolution in time and space of the two environmental variables is defined by the following system of PDEs:

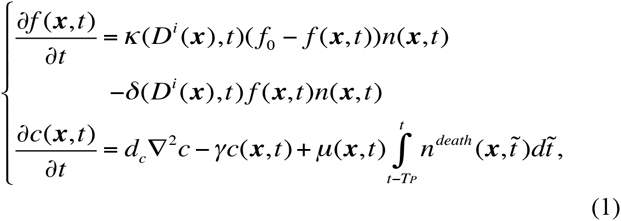

where *d_c_* is the DS diffusion rate, *μ* and *γ* are the constant DS production and decay rates. *κ* and *δ* and are the ECM production and degradation rates, taken as linear functions of the cell phenotype. *f*_0_ is the maximum ECM concentration. Boolean-valued functions n(**x**, t) and n^death^(**x**, t) indicate the presence of a cancer cell or dying cancer cell, respectively. These functions represent the presence or absence of the discrete cancer cell population and will be coupled with a two dimensional numerical discretization of the environmental variables, as described in the supplementary material.

The niche of a cancer cell is therefore defined by the tumor environment, composed of other cancer cells that modulate the production of DS and degrade and move through the ECM. Figure 3 shows a schematic of the interactions between tumor cells and environmental components. An increase in local DS concentration of a cancer cell induces its dedifferentiation, i.e. the acquisition of a more stem-like phenotype, promoting proliferation, and the maintenance of the niche. We define a threshold 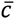 that discriminates between a dedifferentiation-promoting niche 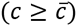, and differentiation-promoting niche 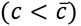. Cell division naturally moves a cell one step towards a more differentiated phenotype.

**Fig. 3.**
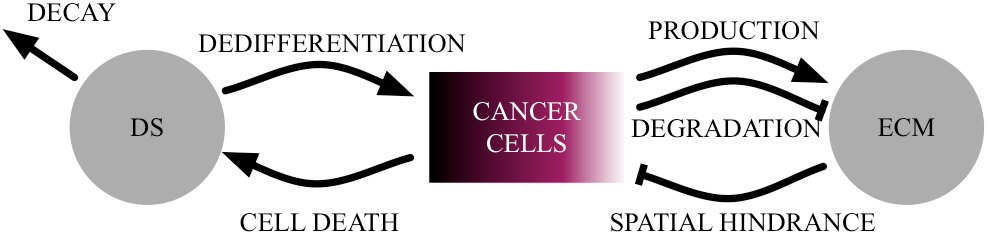
Schematic of stem cell plasticity driven by tumor-environment interactions. Cancer cells interact with two environmental factors: the extracellular matrix (ECM) and the dedifferentiation signal (DS). The DS promotes cell stemness (dedifferentiation to higher stemness phenotypes), is produced by cell death events and degraded naturally. The ECM is degraded and produced by the cells and limits their migration and proliferation.

### A. The Individual Based Model Setup

All the simulations are initialized with a starting configuration of 30 fully stem cancer cells at *D*^0^, randomly spread across the [0. 6 *cm* × 0.6 *cm*] two dimensional domain filled with ECM. A cell’s phenotype is uniquely determined by its *D^i^*, regulating its ability to proliferate and move through the ECM. Cell movement and proliferation are subject to ECM regulation 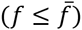. In the context of DCIS, it is necessary to account for the importance of the spatial configuration characterizing the typical ductal structure. Cells that have lost contact with ECM form the inside of a duct, the lumen. If a cell remains in the lumen for too long, this loss of anchorage to ECM, will trigger apoptosis. Finally, fully differentiated cells (*D^N^*), having exhausted their proliferative potential, will be subject to stochastic turnover.

### B. Differentiation and Dedifferentiation Events

The differentiation process corresponds to the natural depletion of a cell’s proliferative potential, due to the progressive telomere shortening at each mitotic event. The remaining proliferative capacity is then modeled as the number of sequential mitoses until full differentiation. A dividing cell at *D^i^* (*i* > 0) will give rise to two daughter cells at *D*^*i*+1^, both with reduced proliferative capacity of (*N* − *D*^*i*+1^). Therefore we can interpret *N* as the maximum number of generations that a cell can give rise to before reaching terminal differentiation. Conversely, a fully stem cell (*D*^0^) can choose between asymmetric and symmetric division. In the first case, it will generate another stem cell (*D*^0^) and a differentiated cell (*D*^1^). In the latter case it will generate two stem cells (*D*^0^). Therefore, by self-renewing, a fully stem phenotype is able to maintain its infinite proliferative capacity. Nevertheless, the modulation of a more or less favorable niche will change the actual proliferative ability of each cell, regardless of its stem state. A favorable niche 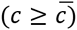 will induce dedifferentiation to *D*^*i*−1^, increasing the cell’s proliferative ability.

Given our definition of niche as the set of environmental conditions allowing cancer cell survival and promoting its stemness, it is possible for the niche to be lost in two different ways. The first is when a cancer cell loses contact with the external matrix, for example in a highly crowded environment, where it is surrounded by its own offspring. The second case is when the local DS concentration falls below 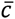 for an extended period of time (longer than the *niche memory*, *T_M_*). In both cases, regardless of the stem state of the cell, the niche is lost and differentiation of one step occurs (from *D^i^* to *D*^*i*+1^). This rule mimics the cancer stem cell dependence on its niche, necessary for the maintenance of the stem-like status.

The HDC approach allows the coupling of discrete events happening at the single cell scale, with continuum environmental variables evolving in space and time at the tissue scale. At each time step all cells are examined, and their fate and phenotype are determined by the current environmental conditions. Subsequently, based on the phenotype, it is determined if proliferation can be carried out. Otherwise, migration is attempted. The collective behavior of the cancer cell population determines how the environmental factor concentrations are modulated (degradation and production of ECM, production of DS). The PDE system is then numerically solved to update the continuum environmental variables. These local updated concentrations then in turn determine the cell’s fate and/or phenotype shift. This whole process is iterated for each simulation time step. A more detailed explanation of the simulation process, model parametrization, and complete flow charts of each process (update of cell fate and phenotype, proliferation and migration) are reported in the supplementary material.

## IV. Results

### A. Recapitulating Stages of Breast Cancer Progression

Since the niche is both modified by the tumor cell and dictates the cell phenotype, we want to investigate both direct manipulation of the niche as well as how the tumor cells perceive it. We therefore consider the impact of varying both tumor and environment properties. Environment wise we consider DS persistence, the ability of the environment to retain the dedifferentiation signal, through delayed degradation or binding to the ECM. The tumor centric properties we consider are the susceptibility of a cell to the DS, i.e. the DS threshold that induces shifts of the cell phenotype, and the time that a cell takes to differentiate once the DS falls below this threshold i.e. the niche memory.

The dedifferentiation signal is an abstract representation of environmental variables that modulate the stemness of cancer cells. The biological counterpart, of this abstract signal, would be more accurately described by an ensemble of many factors (as discussed above) that ultimately determine a cell’s differentiation state. Therefore parameters related to this continuum variable can neither be found in literature nor be measured, but can at least be the subject of a parametric study. By varying values of these three parameters (dedifferentiation signal threshold 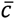, niche memory *T_M_*, and DS persistence *T_P_*) our simple model can produce a suite of different spatial and temporal dynamics. The resulting tumor configurations can be related to those observed clinically at different stages in breast cancer progression, therefore giving an alternate explanation to the variation in pathology across patients. In Fig. 4 we present three simulation outcomes obtained with different values for our three parameters. For all of the simulations presented, the system was initialized with 30 fully stem cancer cells (*D*^0^) randomly distributed in the spatial domain Ω, uniform normalized ECM concentration and zero DS concentration (*f*(***x***, 0) = 1 and *c*(***x***, 0) = 0, ∀ ***x*** ∈ Ω).

**Fig. 4.**
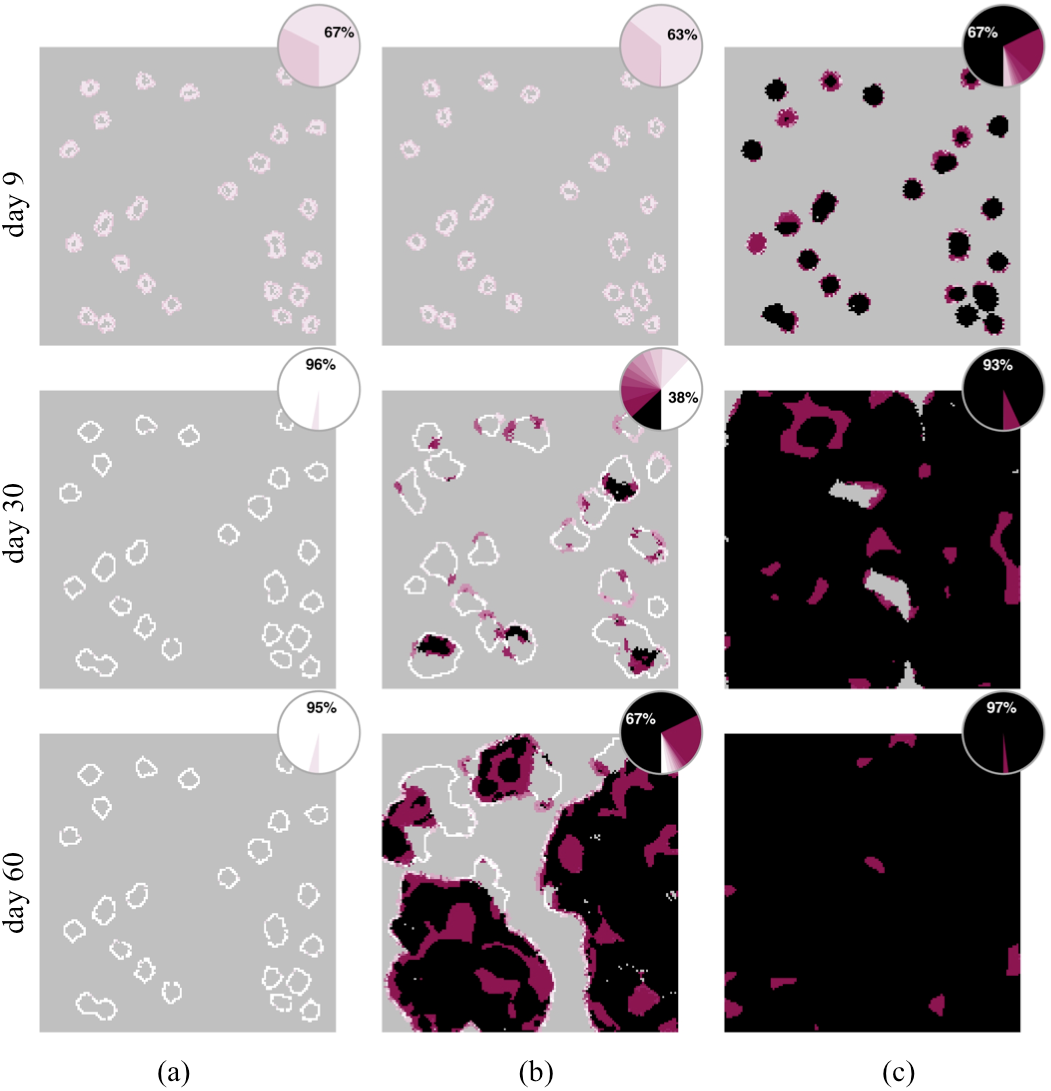
Three representative outcomes obtained varying the DS threshold 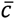, niche memory and DS persistence *T_P_*. In each column three frames of one simulation are shown, after 9, 30, and 60 days of simulation time. Darker shades of purple correspond to higher level of stemness (lower *D^i^*), lighter shades of pink correspond to more differentiated cells (higher *D^i^*), according to the color scheme in Fig. 2. The same color scheme is used in the pie charts, showing distribution of the stem phenotypes. Results span from a well differentiated tissue, with well defined ductal structures which reach a steady state configuration (a), through an intermediate scenario where almost-ductal structures coexist with highly stem-like regions (b), to a scenario where high stemness is the predominant phenotype, leading to a spread invasion of the tissue (c). Insets represent the distribution of *D^i^* for the corresponding frame. Parameter values: 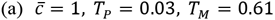; 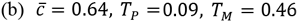; 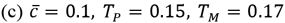. All time constants are measured in days. The square domain is 150 cells wide. Full animations are available in the supplementary material.

In the first simulation [Fig. 4(a)] parameter values are characteristic of an environment unfavorable to the emergence of a stem cell niche, with cancer cells being highly sensitive to differentiation cues. The outcome is a set of well-defined ductal-like structures, emerging from single cancer cells initially seeded in the domain. These ducts are formed by the differentiated progeny, which expands degrading the surrounding matrix. The hollow lumen that these structures display is the consequence of death following crowding and loss of contact with the external ECM. The burst of death occurring in the center of these clusters (Fig. S2) increases the local concentration of DS. However, this does not grow above threshold 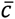, impeding dedifferentiation and re-emergence of stemness. The pie charts show the temporal evolution of the stemness phenotype distribution in each tissue (colors match Fig. 2). Despite starting with fully stem cells, the system gradually progresses towards the fully differentiated phenotype. The white ducts in the last frame are lacking any stem-like trait and represent an equilibrium scenario in which a homeostatic balance has been achieved. This first simulation shows similarities to the structures observed in pathological sections of early breast cancer, i.e. the well-differentiated intra-ductal carcinoma of Fig. 1(a).

In our model a cancer stem cell in the right context can be the source of its own niche: its higher proliferative ability allows it to divide faster and more abundantly. Its numerous progeny will accumulate, dedifferentiate and eventually die, producing high levels of DS that subsequently propagate the niche. The second simulation [Fig. 4(b)] represents an example of this trade-off between the differentiating and dedifferentiating cues. Here, the parameters chosen are more favorable to the accumulation of DS in the environment, while the tumor cells have an intermediate responsiveness to the niche. The outcome is a moderately differentiated tissue, where the different stem phenotypes coexist. Compared to the previous simulation, ductal structures expand more and merge with others in the surrounding tissue. The lower DS threshold allows for the emergence of higher stem phenotypes, while the higher - but not excessive - niche memory allows full stem cancer cells to readily differentiate soon after their appearance. This feedback between cells and environment creates ducts in which distinct regions of differentiation and dedifferentiation appear simultaneously. The pathological counterpart for this moderately differentiated tissue can be seen in Fig. 1(b) where ductal structures are still present, but their lumen are filled.

In the third simulation [Fig. 4(c)] the parameter values characterize an environment in which the emergence and maintenance of the niche is extremely easy. At the same time a cells niche memory is sufficient to delay or escape differentiation and to allow more time for higher DS concentrations to be restored. The outcome is a poorly differentiated and highly stem-like tissue, where phenotypes with higher stemness have taken over the more differentiated ones. Ductal structures that appear in the early stages emerge from a surge of stem proliferation where progeny can easily maintain their stemness. Overall, the cancer stem cell niche easily emerges and is always maintained. Further, the highly stem phenotypes are maintained almost independent of the external environment, and differentiation cues are less effective than in the previous two simulations. In this scenario ductal structures keep enlarging and will eventually merge and fill the entire spatial domain. The tissue is completely dedifferentiated, and resembles the pathology of advanced breast cancer [Fig. 1(c)] where well-defined ductal structures are completely absent.

Corresponding plots of the DS and ECM concentrations for all of the simulations are shown in supplementary material, Fig. S4 and Fig. S5, respectively. It is worthwhile noting that the DS buildup does not decrease monotonically when moving from poorly to highly differentiated tissues. The highest local DS concentration is achieved in the second simulation, corresponding to the intermediate stemness/differentiation outcome. In the case of a more strict niche requirement (high DS threshold and short niche memory), the DS is rarely enough to trigger stemness in cells [Fig. S2(a)]. In the extreme case where niche is easily created and maintained (low DS threshold and long niche memory) the domain is soon filled with highly stem phenotypes and DS builds up rapidly, from dying differentiated cells [Fig. S2(c)]. However, in the intermediate scenario [Fig. S2(b)] the parameter values define a cancer-environment feedback, where alternate waves of niche acquisition and stemness promotion give way to niche loss and differentiation.

### B. Niche Dynamics

In order to better understand the dynamic and emergent nature of the niche, we tracked the values of ECM and DS that all cells experience throughout each simulation to produce an environment-phenotype map. In Fig. 5 (first row) we can identify the clustering of highly stem cells (black and darker shades of purple) and highly differentiated cells (white). For each of the three simulations representing each stage of progression, clusters appear in different areas of the space modulated by the different DS thresholds. We can correlate the highly cancerous stem cell niche with high DS and low to medium ECM. Conversely, the niche of differentiated, DCIS-like cells is characterized by low DS and medium to high ECM. However, it important to note that from a cells perspective the niche does not correspond to a single point in ECM/DS space defined by the model parameters, in fact its far more dynamic and emerges from the collective, plastic, cell-based decisions that occur during a cells lifespan modulated by environmental factors and neighboring cells. Its dynamic nature can be readily appreciated by tracking back the lineage of a few individual cells that are present at the end of a simulation. Representative lineage trajectories for each of the simulations are shown in Fig. 5 (second row). In the intermediate scenario (b, orange/blue lines) lineages can follow very different trajectories as cells that are fully differentiated and dedifferentiated can coexist. However, in the other two scenarios (a, c), all trajectories are similar to the one shown, as cells converge to the same area in the ECM/DS space.

**Fig. 5.**
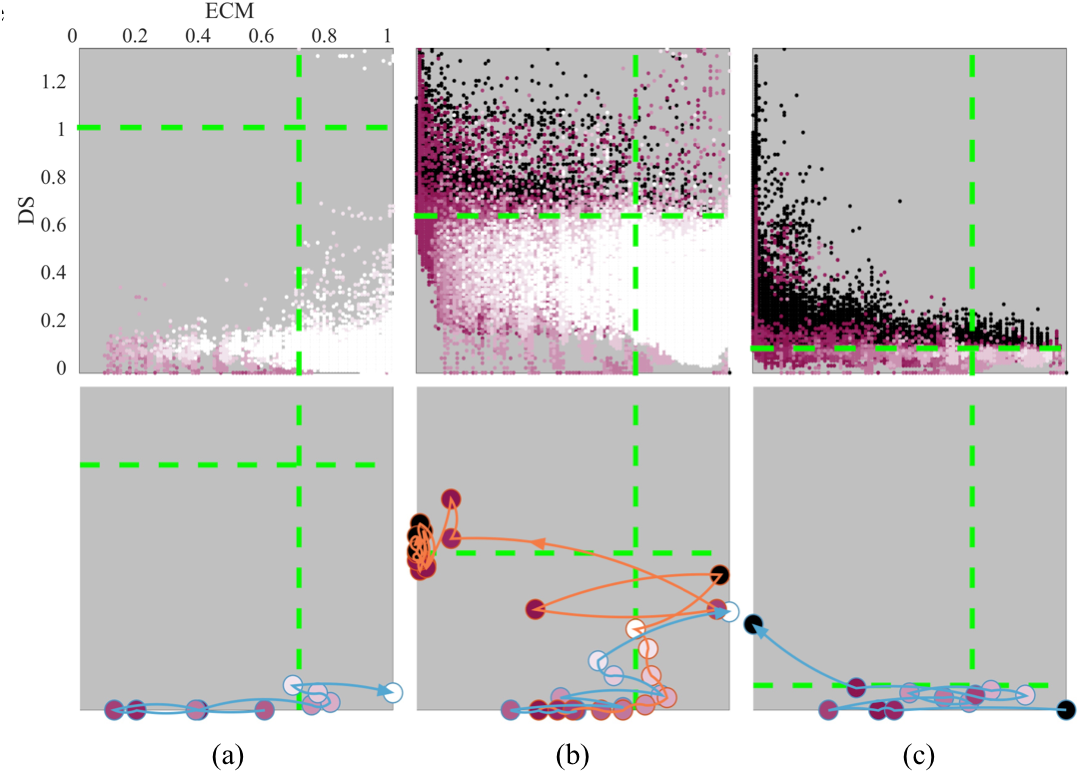
Niche-phenotype map corresponding to the simulations presented in Fig. 4, representative of the range of pathological stages: (a) well differentiated (DCIS), (b) moderately differentiated, (c) poorly differentiated. Color-coded *D^i^* phenotype for all cells against their DS and ECM values for all time steps of the simulation (first row). It is possible to identify regions corresponding to a cancer-promoting niche versus a normalizing one (black vs white areas). Green lines indicate thresholds 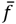 and 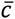. The dynamic nature of the niche can be better appreciated by following the path of a representative single cell lineage trajectory (orange/blue lines) in the DS-ECM space, for each of the corresponding simulations (second row). Color-coded circles along these lines indicate changes in *D^i^* over time.

### C. Distribution of Cancer Cell Generations

Historically there has been some debate as to the number of divisions the progeny of a stem cell is capable of, with numbers varying between 10–20 generations (theoretical range estimated for cancer cells) and up to 50 during development (experimentally estimated for fetal cells) [12], [39]. In our model, even though each mitotic event generates two daughter cells with *D^i^* increased by a unit until full differentiation, the number of grades of stemness *N* does not necessarily correspond to the maximum number of proliferative events a cancer cell can have. Given the environmental modulation of stemness and the cells modulation of the environment, cancer cells are repeatedly subject to dedifferentiation events that confer increased proliferative capacity. The histograms in Fig. 6 show the temporal distribution of generation number for each simulated tissue in Fig. 4. Both the mean and the range vary depending on the parameter set considered, with a tighter mean of 10 generations seen for the stable duct structures in Fig. 6(a) and larger mean and range seen in Fig. 6(b) and (c). When the stemness is highly modulated by the constant feedback between the cancer cells and their niche, cells are more subject to dedifferentiation, which delays their natural progression to full differentiation. Therefore the higher number of generations in Fig. 6(b) is because cells are subject to the highest number of phenotypic shifts. In this parameter regime many cells are likely to undergo the differentiation process in full, and then be consequently shifted to the highest stem-like phenotype, increasing their proliferative potential by *N* generations. This variation in generation numbers (from 10-50), modulated by the niche, may explain why there have been such differences observed in the experimental literature.

**Fig. 6.**
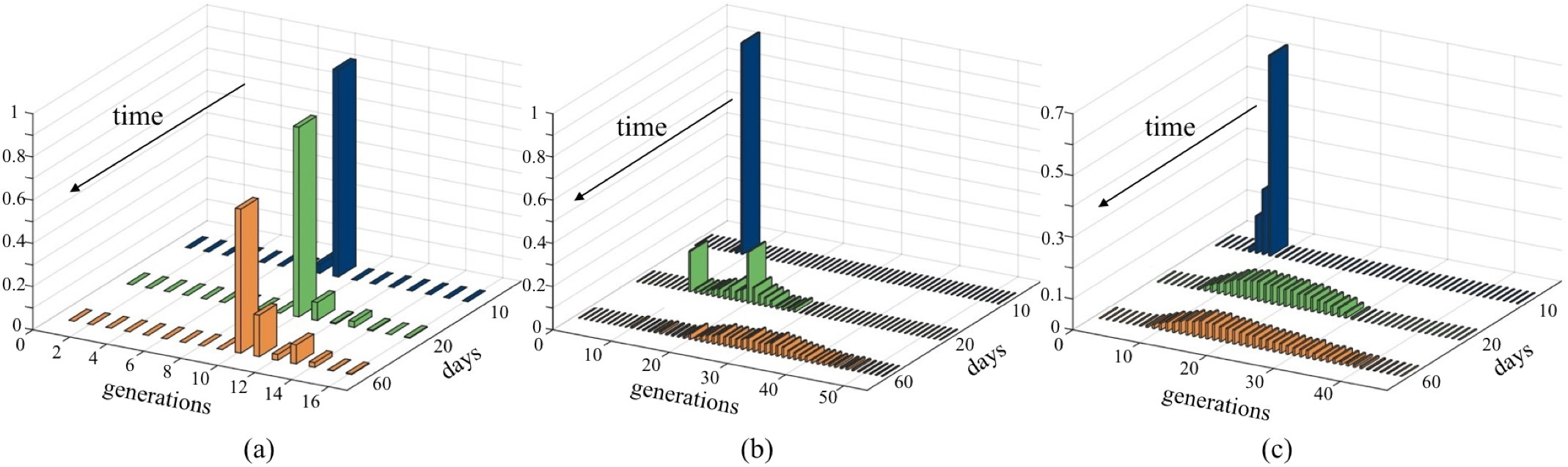
Time evolution of the proliferative ability of cells in the simulations presented in Fig. 4. The distribution of generation number within the cancer cell population at a given time point is normalized to the total population number at that time. (a) In the highly differentiated scenario cancer cells are very unlikely to undergo dedifferentiation, hence most cells possess a proliferative ability equal to *N*, the number of generations to full differentiation. (b) In the intermediately differentiated scenario, the stem phenotype is more modulated than in the previous case. Cancer cells may go through all the *N* differentiation steps before their phenotype is shifted back to the highest stem-like one. (c) In the poorly differentiated scenario, stemness is highly maintained, allowing the cells to increase their proliferative potential by tens of generations.

### D. Exploring the Parameter Space

It is possible to explore the full three dimensional parametric space to observe the distribution of scenarios when cell-centric (dedifferentiation signal threshold 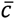 and niche memory *T_M_*), and environment-centric (DS persistence *T_P_*) properties are varied. In order to obtain an informative representation of the 3D parametric space, we classify each simulation outcome as either: poorly, moderate or highly differentiated, as referred to in the equivalent clinical counterpart (Fig. 1). Figure 7 shows this classification for a complete set of simulations spanning the parametric space within interesting and biologically realistic value ranges (details in the supplementary material). Consistent with the previous results, highly differentiated tissues appear in the high 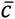 - low *T_M_* - low *T_P_* region, where niche acquisition is difficult. Conversely, highly invasive and dedifferentiated tissues result in the stricter parameter regime (low 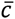 - high *T_M_* - high *T_P_*). The transition between these extreme scenarios is non linear and the shape of this parameter space would benefit from further investigation. The full set of simulations is available in the supplementary material.

**Fig. 7.**
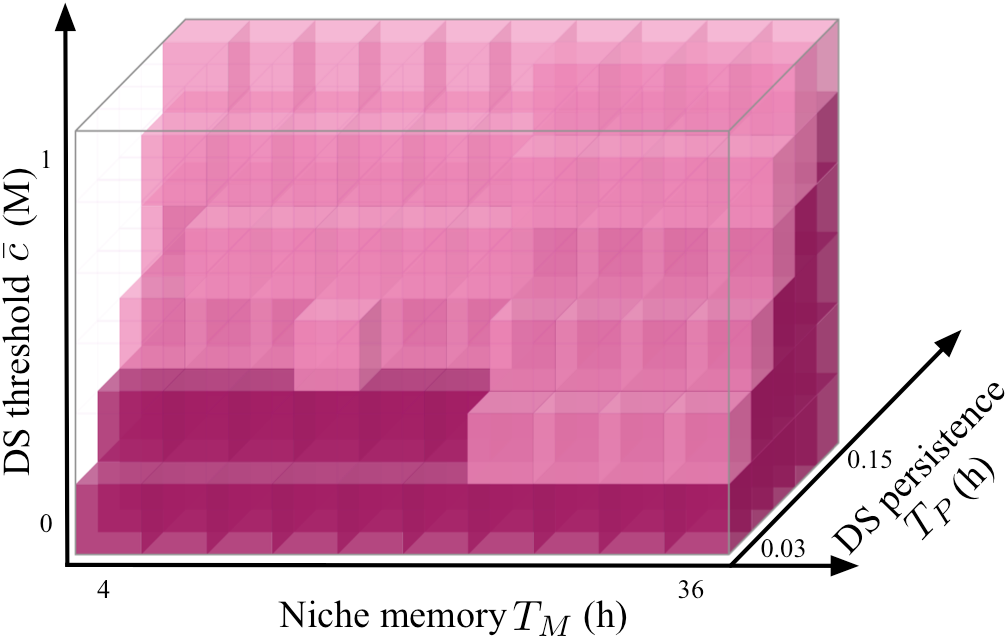
Exploration of the 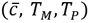 parameter space. Simulation outcomes are classified as either: *poorly differentiated* cancer if all cells have reached and maintained highly stem phenotypes (purple), *highly differentiated* cancer if all cells are fully differentiated (transparent), *moderately differentiated* cancer for the other intermediate cases, where highly stem and highly differentiated phenotypes coexist (pink). Classification is made at time 60 days, when most of the simulations have reached confluence or convergence.

### E. Invading Homeostatic Ducts

So far we have explored a range of scenarios that can emerge from single cancer cells that are able to, more or less, recapitulate the ductal structures of the primary breast tissue. We now explore the model dynamics from the point of view of tumor initiation in pre-existing homeostatic ductal structures. Specifically, we consider the impact of different mutations affecting an individual cell lining one of the ducts. The position of such a mutant cell within a differentiated duct is the same in all of the following simulations. The homeostatic tissue in which we will seed the mutants is obtained from the previous simulations. Specifically, we choose one of the well differentiated structures and we initially suppress cellular turnover, in order to recapitulate a well established and robust structure.

The first type of mutation considered is one altering the cell’s response to the environment. Namely, its niche memory *T_M_* is infinite and its dedifferentiation signal threshold 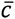 is zero, meaning that the mutant cell is completely unresponsive to environmental and differentiating cues. This is the case of a cell hijacking the regulatory mechanisms in the niche that previously allowed the formation of differentiated structures. In Fig. 8(a) three snapshots of this simulation are shown. Despite the normalized, homeostatic tissue in which the mutation is introduced, the mutant cell easily acquires highly stem-like properties and, being unresponsive to the environmental signals, maintains such phenotype. The mutant progeny easily invade the surrounding space. However, the growth is limited to the lumen, similar to DCIS where cancerous growth is limited by an intact basement membrane. If we were to introduce the very same mutant cell within a less stable structure, i.e. where turnover of fully differentiated cells is allowed, we would initially obtain similar dynamics [Fig. 8(b)]. However, after filling the lumen, taking advantage of the differentiated cells turnover, the invasive tumor mass breaches the duct into the surrounding space. These two examples suggest that mutations that render cells irresponsive to the regulatory mechanisms of a tissue are not sufficient to induce invasive growth. Indeed, as long as the structure in which they arise is stable enough, i.e. a non cancerous breast duct like in Fig. 8(a), the effect of its growth is limited. Conversely, if the structure is somehow destabilized (in this case by turnover of cells lining the duct), the mutant is able to break through and form one big expanding mass. However, the mutant lineage does not impact how the environment modulates the other cells, which remain stable hollow ducts.

**Fig. 8.**
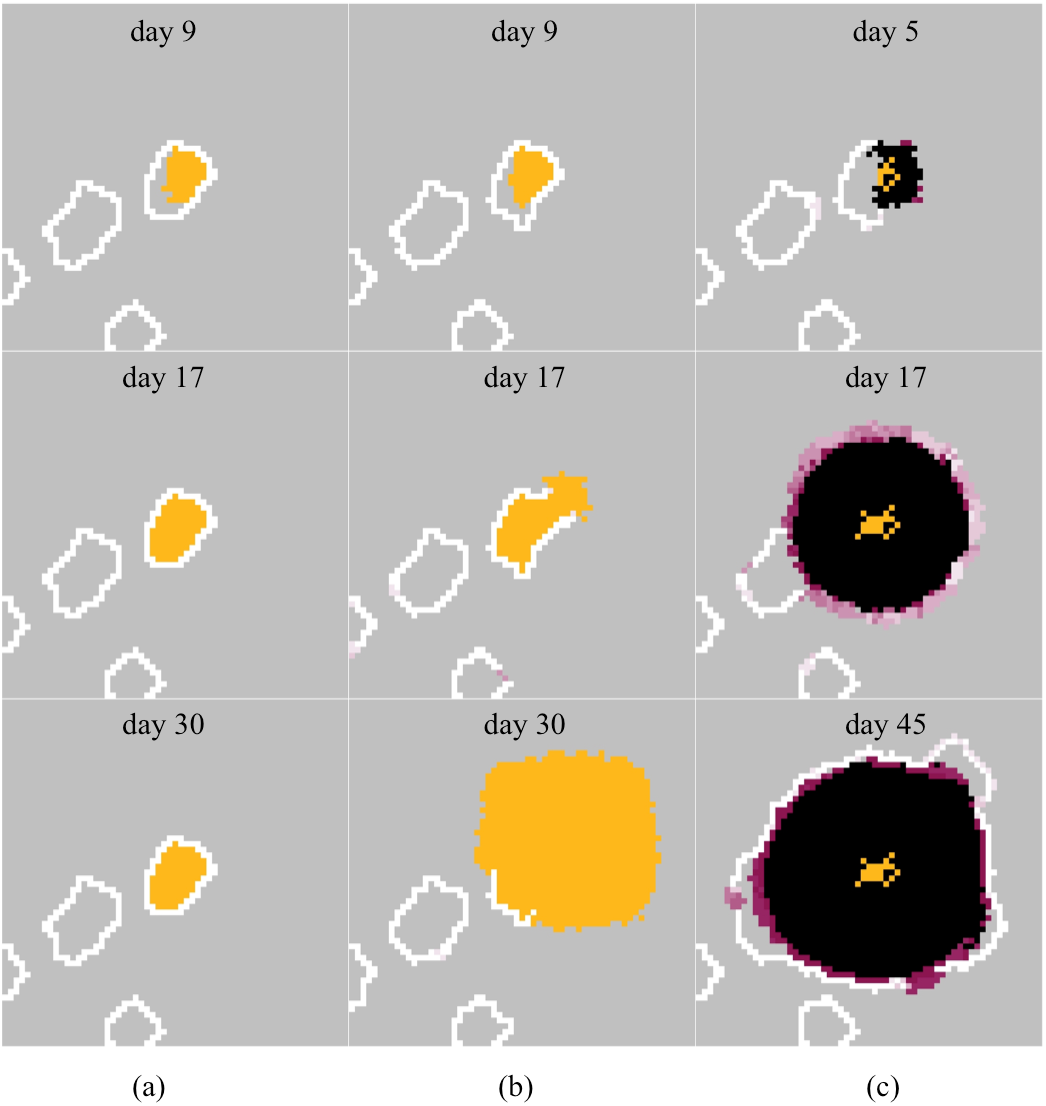
A mutation is introduced in a pre-existing homeostatic tissue with fully differentiated ducts. Three frames of each simulation are shown, at indicated days from the mutation time. The mutant cell and its progeny are tagged in yellow. The homeostatic tissue is characterized by 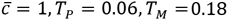. Other parameter values are the same as in the previous simulations. (a) The mutated cell is characterized by *T_M_* = ∞ and 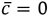, making the cell completely irresponsive to the environmental cues. The mutation is introduced in a non-cancerous tissue with stable ducts. (b) The same mutation is introduced in a well differentiated cancerous tissue, characterized by turnover. The mutant’s progeny breaches the duct. (c) A different type of mutation is introduced in a well differentiated cancerous tissue. The mutated cell actively modifies the environment through extra production of DS, causing a bystander effect and invasive growth.

The third type of mutation we consider [Fig. 8(c)] is one that causes the cell to actively modify its environment by upregulating the production of DS. The modified equation is reported in the supplementary material. Interestingly, the feedback between the “normal” behavior of the pre-existing differentiated ducts and the environment altered by the mutant cell and its progeny drives somewhat different dynamics. The buildup of DS, emanating from the mutant cells and diffusing into the surrounding tissue, creates an ever expanding niche which induces dedifferentiation in the pre-existing fully differentiated cells propagating the large highly stem core. This effectively represents a bystander effect where, unlike the previous cases, the mass is mostly made of the progeny of non-mutant cells. It is worth noting that the results bare some resemblance to those of Fig. 4(b), where we observed a tradeoff between differentiation and dedifferentiation.

## V. Discussion

Our model explores the hypothesis that the cancer cell environment is a major driver of the cell phenotype regulating stemness. unlike many models of cancer stem cells it considers a continuous gradation of stem like phenotypes, challenging the hierarchical model of stemness.

Starting from a baseline ability to generate homeostatic normal ducts, we demonstrate that the model can recapitulate the dynamics of early invasion and progression in primary tissue. Modifying a few parameters that relate to niche acquisition and maintenance, we find that the model can capture the different stages of ductal cancer progression (compare Fig. 1 and Fig. 4). In addition, we observed that the stem cell niche is a dynamic, both spatially and temporally, emerging property of the system (Fig. 5). Alternatively, by introducing a mutation, which causes a cell to shift its environment toward a more stem-promoting niche, we observe an invasive phenotype that breaches the duct structure, invading out of the lumen and into surrounding tissue (Fig. 8). Furthermore the model predicts that this mutant also induces dedifferentiation in any pre-existing differentiated cell in the surrounding ducts through manipulation of the niche.

The role of CSCs is profoundly important for understanding the dynamics of tumor growth because it ultimately defines the “unit of selection” in somatic evolution. In other words, what organism is being selected during somatic evolution of cancer? Most investigators would identify the individual cancer cell as the unit of selection in cancer biology. Thus, the Darwinian dynamics will select for properties in individual cells that maximize proliferation within the context of the local environment. However, if the hierarchical model of CSCs is correct, then the unit of selection must be the multicellular structure that consists of the stem cell and all of its progeny. In this model, cancers are composed of self-replicating modules that, similar to bee hives, consist of 100s or 1000s of cells. Thus, natural selection would optimize a module configuration that maximizes the self-renewal of each stem cell that would then form another module. In contrast, if the unit of selection is individual cancer cells, then it would seem most likely that the “stem phenotype” is part of the cellular reaction norm i.e. a cell property that could be gained or lost depending on environmental cues, as we propose here.

Normal somatic stem cells and their niche are fundamental units of normal tissue development and homeostatic regulation. It would, therefore, not be surprising for tumor populations to have a similar organization. This hierarchical model of CSCs proposes that, similar to normal tissue, cancer stem cells give rise to an ensemble of more differentiated cells that, similar to a beehive, for example, maintain favorable environmental conditions (i.e. the stem cell niche) that promotes self-renewal of the stem cells. However, we note that such an arrangement requires each stem cell to produce 100s or 1000s of cells that cannot self-renew and therefore do not directly contribute to the clonal line. While certainly possible, we note that no organized structures of such magnitude are reported in cancers. Furthermore, the production of multiple, genetically identical copies of a stem cell (but without the capacity to proliferate indefinitely) represents a large expenditure of resources for no apparent evolutionary advantage. Thus, we explore an alternative model in which the Darwinian unit of selection is an individual but highly plastic cancer cell. In this model, feedback between the environment and the tumor cell may result in a stem phenotype, which need not be permanent [5], [40]. Therefore stemness can be acquired and lost purely dependent on environmental context. Among the numerous environmental factors thought to play a role in this process are cellular and soluble components, signaling molecules and stress factors released under harsh environmental conditions. These factors contribute to the modulation of the cancer stem cell niche and have been identified as drivers of dedifferentiation, either inducing or maintaining the stem cell phenotype [33], [41].

## VI. Conclusion

To investigate the environmentally-driven stem cell plasticity hypothesis, we developed a Hybrid Discrete Continuum model of DCIS. This allowed us to carry out an *in silico* investigation of the dynamics occurring during progression at the single cell scale in dialogue with the tumor microenvironment.

Our results suggest that for ductal cancers, the mechanisms that ensure normal duct formation and maintenance are the very ones that cancer inadvertently hijacks by becoming a niche manipulating machine. Cancers ability to modify its environment has been known for sometime and has even prompted some novel therapeutic approaches (e.g. anti-angiogenic and anti-protease therapies) with limited success. One issue these approaches ignore, that our work emphasizes, is that the niche is a dynamic and emergent property of the collective interactions between cells and environmental factors. Therefore the niche is a moving target that can’t be controlled by only targeting one of these factors. Instead, we need to develop therapeutic strategies that target both the tumor and its environment, such that normal homeostatic regulation can be reinstated.

In conclusion the central idea, that plasticity of the stem phenotype is largely context driven (i.e. part of the cellular reaction norm) provides an alternate interpretation of the initiation and subsequent progression of ductal carcinoma, since it naturally captures the spectrum of dedifferentiation observed clinically. We also hope our results motivate a different therapeutic perspective that explicitly accounts for the dynamic and emergent nature of the niche. Finally, we acknowledge that our results are significantly dependent on the ability to both define and quantify the impact of environmental signals which drive dedifferentiation – a difficult task since such signals are poorly studied. However, there is some intriguing experimental evidence that hints at the existence of such a signal, but gives no explicit qualification of its impact in terms of stemness promotion [28], [40].

## Acknowledgment

The authors would like to thank Moffitt investigators, Drs. Heiko Enderling and Jacob Scott for useful discussions during the development of this work. We would also like to thank Prof. Luigi Preziosi for initiating this collaboration.

